# ACIDIC BICELLES ARE A SUITABLE MEMBRANE MIMIC FOR STRUCTURAL STUDIES OF THE LASSA VIRUS FUSION DOMAIN

**DOI:** 10.1101/2025.03.11.642635

**Authors:** Hallie N. Pennington, Jinwoo Lee

## Abstract

Lassa virus (LASV) is the most prevalent arenavirus afflicting humans and has high pandemic potential. The genetic material of LASV is delivered into the host cell via membrane fusion – a process initiated by the LASV fusion domain (FD). However, the molecular details of the LASV FD, particularly its structure after association with the host cell, remain poorly understood. This can be attributed to a lack of a viable membrane mimic to effectively stabilize the LASV FD for structural studies. Here, we demonstrate that the structure of the LASV FD widely varies based on the class of membrane mimic. In particular, through CD spectroscopy, we found that the LASV FD required a charged membrane mimic, such as zwitterionic or anionic detergent micelles, to adopt a helical conformation at low pH, but has the highest helical content in the presence of anionic lipids, particularly the detergent micelle LMPG and acidic bicelles. Moreover, we reveal that the LASV FD was well resolved on NMR spectra in CHAPS, DPC, LDAO, LMPG, and acidic bicelles, where LMPG and acidic bicelles had the sharpest peak resolution, but more defined peaks were noted in acidic bicelles over LMPG. In conclusion, our findings indicate that acidic bicelles are the optimal membrane mimic for the stabilization of the LASV FD such that structural studies can be conducted.

**Graphical Abstract:** 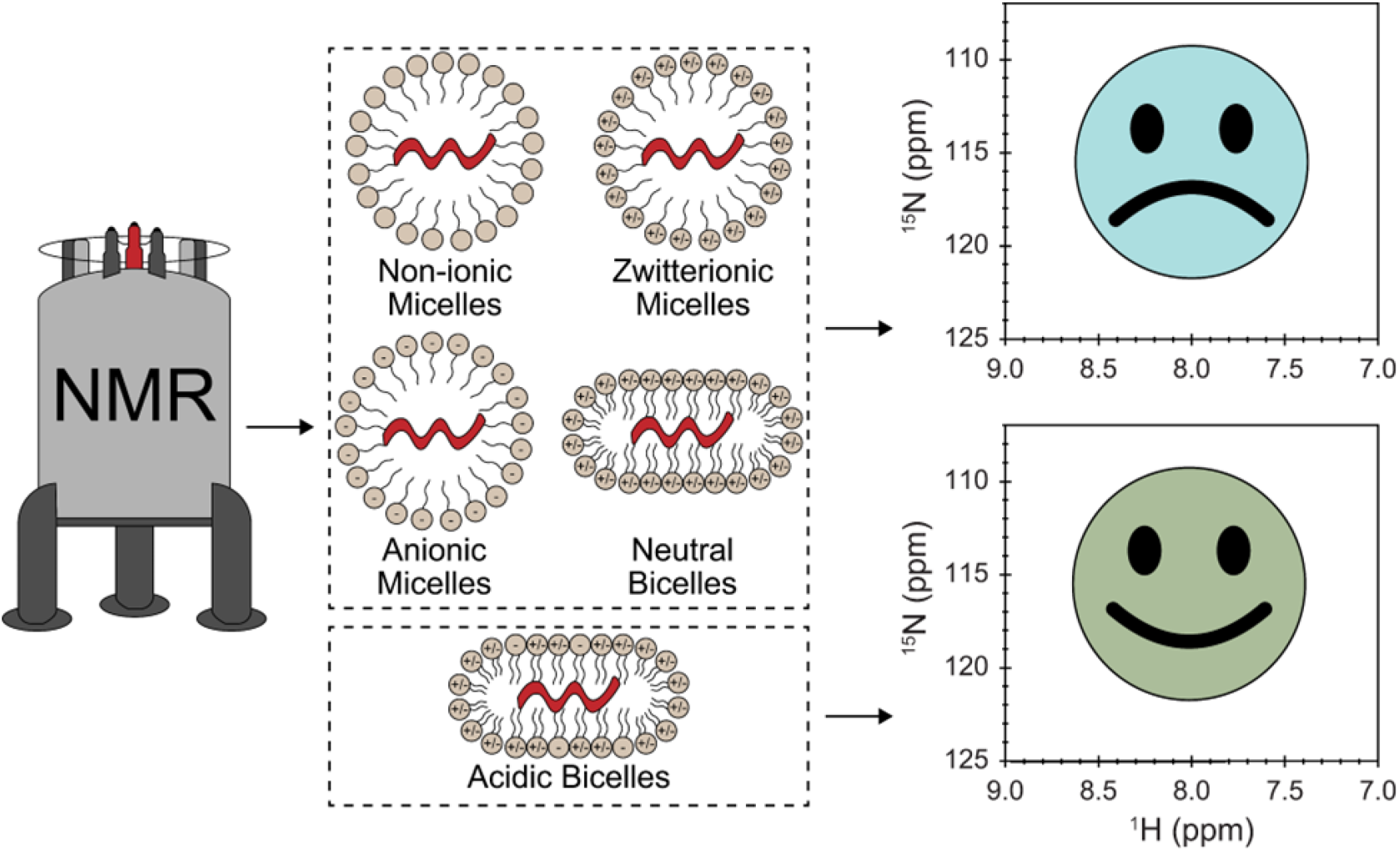

## Introduction

Infection with Lassa virus (LASV), an emerging arenavirus endemic to West Africa, causes Lassa fever – a lethal hemorrhagic fever^1-3^. On an annual basis, 100,000 to 300,000 individuals are infected with LASV, but these numbers may be dwarfed due to poor healthcare infrastructure in endemic regions^4-7^. Unfortunately, up to 20% of those infected with LASV will ultimately succumb to the virus, particularly in severe cases. LASV is most commonly spread by direct contact with infected rodent excrement, but an uptick in direct, human-to-human transmission has been noted in recent years. Despite the evolving pathogenicity of LASV, there are presently no FDA-approved vaccines or antivirals^8-11^. Thus, the World Health Organization (WHO) has deemed LASV as one of the top five diseases requiring prioritized research and development due to its pandemic potential as it nears a complete zoonotic jump^12^.

LASV’s genetic material is delivered into the host cell via membrane fusion – a process mediated by the glycoprotein complex (GPC), which is the sole protein located on the virion surface^13-15^. The GPC is comprised of three subunits, including a receptor binding subunit (glycoprotein 1, GP1) and a membrane fusion subunit (glycoprotein 2, GP2) that is associated with a stable signal peptide (SSP). After an initial interaction between GP1 and its primary receptor on the host cell plasma membrane (α-dystroglycan (αDG)), the virion is endocytosed and delivered late in the endocytic pathway where fusion ultimately occurs [Figure 1]. It is believed that LASV-mediated membrane fusion occurs via the six-helix bundle mechanism (6HB), similar to the process undergone by other class one fusion proteins like severe acute respiratory syndrome coronavirus 2 (SARS-CoV-2), influenza, human immunodeficiency virus (HIV), and Ebola virus (EBOV). Thus, the initial step of LASV fusion likely involves the anchoring of a hydrophobic sequence at the N-terminus of GP2, known as the fusion domain (FD), into the host cell membrane. The viral and host cell membranes are then tethered together such that membrane fusion can transpire, resulting in the formation of a fusion pore and merging of the membranes. In this article, we will use the terminology ‘pre-fusion’ and ‘post-fusion’ to refer to the state of the FD before and after associating with the target host cell membrane, respectively.

**Figure 1.**
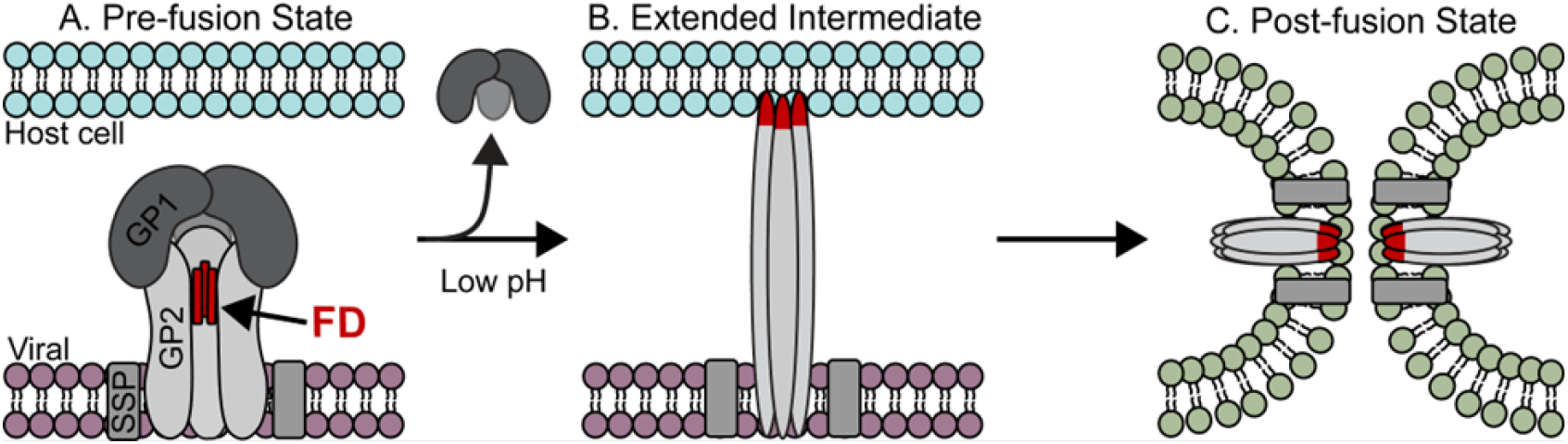
Illustration of LASV viral entry into the host cell via the endocytic pathway and six helix bundle (6HB) mechanism. [A] In the pre-fusion state, the receptor binding subunit (glycoprotein 1, GPI) sits atop the membrane fusion subunit (glycoprotein 2, GP2), which is associated with a stable signal peptide (SSP)-[B] Upon delivery of the virion to the lysosomal compartment, the low pH results in the dissociation of GPI and subsequent exposure of the fusion domain (FD) at the N-terminus of GP2, which embeds itself into the host cell membrane, creating an extended intermediate structure. [C] The host cell and viral membranes are tethered together such that when GP2 folds back on itself, a fusion pore is formed, and the two membranes are merged.

While FDs are generally well conserved in a single viral family, they can vary greatly across different viral families, not only in sequence, but in structure as well. For example, although the FD of both HIV and influenza contain an N-terminal fusion peptide (FP), their post-fusion structures are significantly different with HIV adopting an extended helix, whereas influenza adopts a helical hairpin, boomerang structure^16-19^. On the other hand, the EBOV FD is comprised of an internal fusion loop (FL) that is held together by a disulfide bond and forms a hydrophobic fist at a low pH to mediate fusion^20, 21^. The LASV FD and other arenaviruses are unique in that they contain both an FP (G^260^ – T^274^) and FL (C^279^ – N^295^) that are conjoined by a short linker region (P^275^ – Y^278^)^22^. This feature is shared only with coronaviruses, such as SARS-CoV-2, where the FP and FL are functionally distinct, but are most effective in synergy with the FP adopting a helix-turn-helix motif and FL serving as a mechanical stabilizer^23-28^. While previous work has elucidated the structure of the LASV FD in its pre-fusion state, there has yet to be a detailed investigation into its post-fusion structure^14, 15^.

Previously, we demonstrated that the LASV FD adopts a fusogenic, helical conformation at a pH akin to the lysosomal compartment^22^. However, the location of any helices within the post-fusion state of the LASV FD was not indicated. Additionally, this was accomplished in vesicles, which are unfavorable for structural studies. Given its role in fusion and relative conservation, the FD serves as an important target for the development of novel, broadly neutralizing antibodies and vaccines. This is the case for HIV where a multitude of antibodies have been shown to prevent infection by binding to the FP^29-31^. Antibodies with action against the FD of other viruses, such as influenza, EBOV, and SARS-CoV-2, have been described, as well^32-38^. The regions of these viral FDs that the antibodies target have often been revealed via structural investigation. Therefore, it is pertinent to determine a viable membrane mimic for the LASV FD such that the post-fusion structure can be elucidated to understand LASV FD-initiated fusion and provide insights into regions of potential therapeutic target.

In this study, we employed a detergent screen of nine membrane mimics to deduce the optimal membrane mimic for the stability of the LASV FD. Through CD spectroscopy, we revealed that, at a low pH, the FD transitioned to a helical conformation in zwitterionic and anionic detergent micelles and acidic bicelles but remained in a random coil conformation in non-ionic detergent micelles and neutral bicelles. Collectively, we provide evidence that acidic bicelles are the optimal membrane mimic for structural investigation of the LASV FD.

## Materials and Methods

### Detergents and Lipids

N-lauroylsarcosine (sarkosyl) and N,N-dimethyl-1-dodecylamine N-oxide (LDAO) were both purchased from Fisher Scientific. Dodecyl-β-D-maltoside (DDM) was purchased from Sigma Aldrich. Dodecylphosphocholine (DPC), 3-[(3-cholamidopropyl)-dimethylammonio]-1-propane sulfonate (CHAPS), n-octyl-β-d-glucopyranoside (β-OG), and 1,2-dimyristoyl-sn-glycero-3-[phospho-rac-(1-glycerol)] (DMPG) were purchased from Anatrace. 1-myristoyl-2-hydroxy-sn-glycero-3-phospho-(1′-rac-glycerol) (LMPG), 16:0–18:1 1-palmitoyl-2-oleoyl-glycero-3-phosphocholine (POPC), 16:0–18:1 1-palmitoyl-2-oleoyl-*sn*-glycero-3-phospho-[1′-rac-glycerol] (POPG), 1,2-dimyristoyl-sn-glycero-3-phosphocholine (DMPC), and 1,2-dihexanoyl-sn-glycero-3-phosphocholine (DHPC), were all purchased from Avanti Polar Lipids.

### Expression and purification

The Lassa virus fusion domain (LASV FD) construct ^260^(GTFTWTLSDSEGKDTPGGYCLTRWMLIEAELKCFGN)^295^ was designed with an N-terminal 9x-His tag followed by a trp operon leader sequence (TrpLE), and thrombin cleavage site. The natural thrombin cleavage site was replaced with LVPR↓GT to yield the native sequence of the LASV FD after cleavage. The expression and purification protocol for the LASV FD has been described in detail previously^22^. Briefly, Ni-NTA affinity chromatography and thrombin cleavage were employed to separate the protein from the tags. Size exclusion chromatography (SEC) was then utilized to further purify and isolate the protein. For isotopic labeling of the LASV FD with ^15^N, cells were grown in 1 L EMBL minimal media containing 8 g/L Na_2_HPO_4_, 2 g/L KH_2_PO_4_, 0.5 g/L NaCl, 1 g/L ^15^NH_4_Cl, 10 g/L glucose, 1 mM MgSO_4_, 0.3 mM Na_2_SO_4_, 0.3 mM CaCl_2_, 50 μg/mL kanamycin, 34 μg/mL chloramphenicol, and trace amounts of biotin and thiamine. A 5 mL starter culture in the minimal medium was grown overnight at 37 °C and 225 rpm. The starter culture was propagated into 30 mL of fresh minimal media, grown overnight at 37 °C and 225 rpm, then propagated into 120 mL of fresh minimal media and grown overnight again, before being added to a 1 L volume. The expression and purification of the isotopically labeled FD then followed the same process as described previously_22_.

### Preparation of Unilamellar Vesicles

Small unilamellar vesicles (SUVs) were prepared by mixing appropriate amounts of stock lipid solutions in glass test tubes. A gentle, continuous stream of nitrogen was then applied while vortexing to remove the chloroform and create a lipid film. This film was left in a vacuum desiccator overnight to fully remove any residual chloroform. The next day, after resuspending the lipid film in the appropriate volume of CD buffer (1 mM HEPES/MES/Sodium Acetate (HMA), 10 mM NaCl, pH 7.4) by vortexing, the lipids were sonicated on ice for 15 min (1 s on, 1 s off, 10% power) using a Branson Digital Sonifier SFX 250 equipped with a titanium microtip. The transparent solution was subsequently centrifuged at 20,000 x g for 15 min in an Eppendorf 5424 microcentrifuge. Once any particulates were removed, the lipids were transferred to a fresh microcentrifuge tube for usage. The vesicles utilized in this study were comprised of 65:35 POPC:POPG to provide a rudimentary mimic of the membrane composition found in the lysosomal compartment^22^.

### Circular Dichroism (CD) Spectroscopy

CD measurements were performed on a Jasco J-810 spectrophotometer with a 2 mm quartz cuvette. All experiments were conducted at room temperature (∼22 °C) in 1 mM HMA, 10 mM NaCl, pH 7.4 or 4.0 with a protein concentration of ∼ 10 μM. The concentration of bicelles was 1% with a q-value of 0.5 with either DMPC:DHPC (neutral bicelles) or (75:25 DMPC:DMPG):DHPC (acidic bicelles). At a minimum, all detergents were at a concentration of 10X their critical micelle concentration (CMC), specifically 10 mM LMPG, 20 mM of either DDM, LDAO, or DPC, 100 mM CHAPS, 150 mM Sarkosyl, or 200 mM β-OG. The spectra for the sample in the various membrane mimics were collected at pH 7.4, then the pH was dropped with 1 M HCl to 4.0 as confirmed with a pH probe. The sample was incubated overnight at 4 °C to give the LASV FD enough time to appropriately associate with the mimic before repeating the spectra collection the following day. Samples were spun down for 10 min at 20,000 x g before each spectra collection to deduce any precipitation. Spectra were collected from 260 nm to 198 nm with a step size of 1 nm at 20 nm/min and averaged over three accumulations. Measurements were taken of the buffer alone containing each of the membrane mimics from 260 nm to 198 nm with a step size of 1 nm at 50 nm/min, averaged over three accumulations, and then subtracted from the corresponding sample. Subtractions and smoothing were performed using CDToolX and the helical percentage was calculated as described previously^22, 39^.

### Nuclear Magnetic Resonance (NMR) experiments

NMR spectra were acquired using a Shigemi NMR tube with a sample concentration of ∼300 μM in a sample volume of 300 μL in 25 mM Na_2_HPO_4_, 100 mM NaCl, pH 7.0 or pH 4.0 with a 90:10 ratio of H_2_O:D_2_O. A bicelle concentration of 25% was utilized to maintain the same bicelle:protein ratio as the CD experiments, whereas all detergents were at the same concentration as the CD experiments since they were significantly above their CMC. All experiments were carried out on a Bruker Ultrashield^TM^ 600 MHz magnet with a CP2.1 TCI 600S3 H&F/C/N-D-05 Z XT Cryoprobe at a temperature of 37 °C. All data was processed using NMRPipe and NMRFAM-SPARKY via NMRBox^40-42^.

## Results

### A charged membrane mimic is required for the LASV FD to undergo a conformational change at low pH

To determine the optimal detergent for the LASV FD, we utilized CD spectroscopy to screen a multitude of membrane mimics for stability and secondary structure of the LASV FD at pH 4.0 [Figure 1,2]. This included the detergent micelles dodecyl-β-D-maltoside (DDM), n-octyl-β-d-glucopyranoside (β-OG), N,N-dimethyl-1-dodecylamine N-oxide (LDAO), 3-[(3-cholamidopropyl)-dimethylammonio]-1-propane sulfonate (CHAPS), dodecylphosphocholine (DPC), N-lauroylsarcosine (sarkosyl), and 1-myristoyl-2-hydroxy-sn-glycero-3-phospho-(1′-rac-glycerol) (LMPG). Additionally, we examined the properties of the LASV FD in neutral bicelles comprised of 1,2-dimyristoyl-sn-glycero-3-phosphocholine (DMPC) and 1,2-dihexanoyl-sn-glycero-3-phosphocholine (DHPC) or acidic bicelles comprised of DMPC, 1,2-dimyristoyl-sn-glycero-3-[phospho-rac-(1-glycerol)] (DMPG), and DHPC. These detergents were selected as they are all commonly employed for biochemical or biophysical characterization of various membrane proteins, particularly in solution NMR spectroscopy^24, 43-51^. We found that Sarkosyl was not stable at pH 4.0 as a large precipitate was observed immediately upon acidification [Figure 2A]. All other membrane mimics were stable for a minimum of seven days at pH 4.0 [Figure 2B]. However, despite the stability of CHAPS at pH 4.0, as noted by the lack of precipitation after overnight incubation, the spectrum was extremely noisy despite the removal of the background signal [Figure 3A]. This left us to investigate the structure of the LASV FD in SUVs versus in the detergent micelles DDM, β-OG, LDAO, DPC, and LMPG as well as neutral bicelles and acidic bicelles.

**Figure 2.**
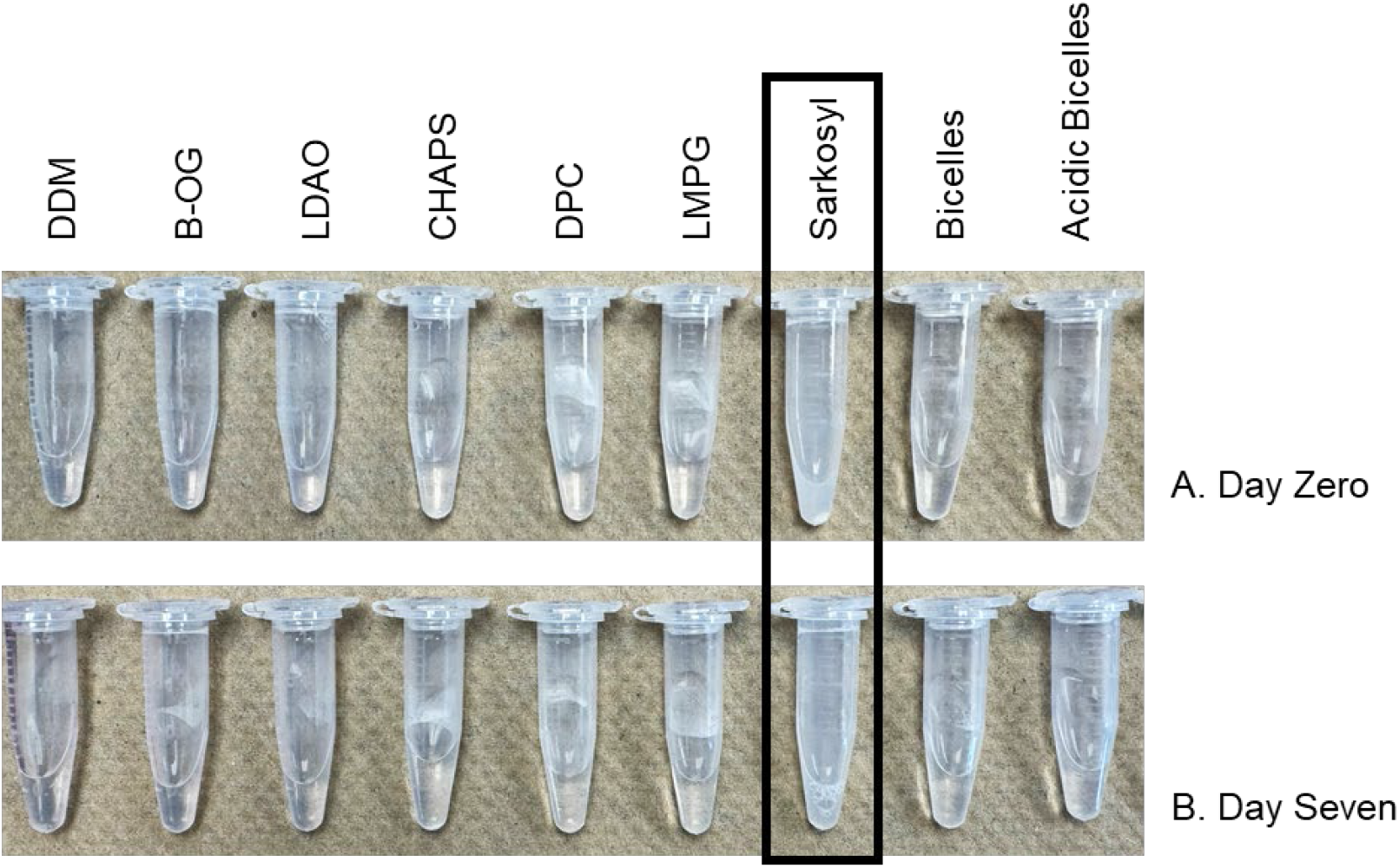
The LASV FD is stable in all the tested membrane mimics, except Sarkosyl (black box), at pH 4.0 fora minimum of seven days.

**Figure 3.**
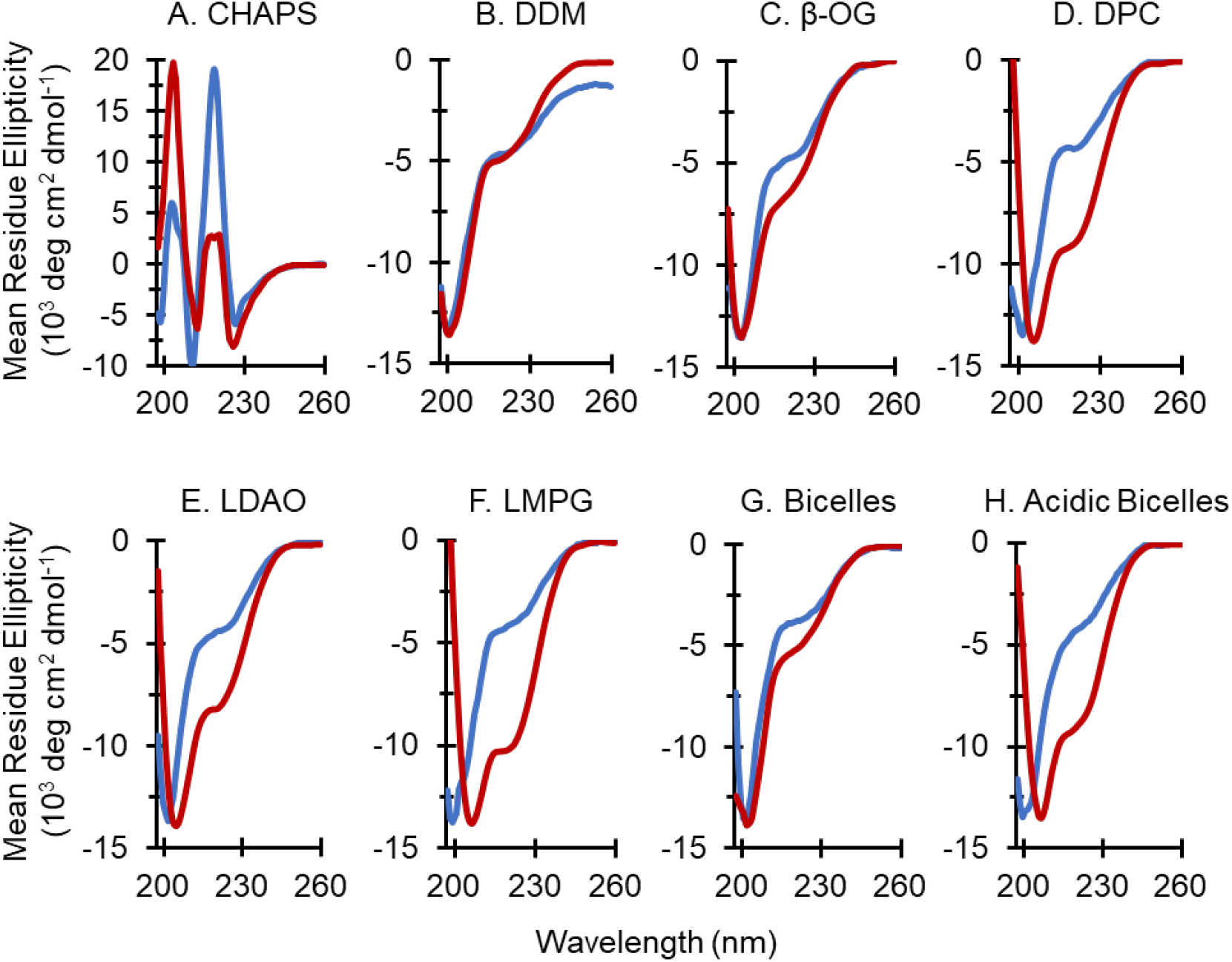
The secondary structure of the LASV FD has a large dependence on the membrane mimic present. The signal-to-noise of the LASV FD in [A] CHAPS was too high to determine the structure at pH 7.4 (blue) or pH 4.0 (red). In the non-ionic detergents [B] DDM and [C] ß-OG, the LASV FD remained in a random coil conformation regardless of physiological pH or low PH. A conformational change of the LASV FD from random coil at pH 7.4 to helical at pH 4.0 was observed in [D] LDAO and [E] DPC as well as [F] LMPG. In [G] neutral bicelles, the LASV FD remained as a random coil regardless of pH but adopted a helical structure in [H] acidic bicelles.

In SUVs, the LASV FD was stable with a random coil conformation at physiological pH (7.4) and had a helical conformation at low pH (4.0)^22^. While the LASV FD had a similar, random coil structure in all membrane mimics at physiological pH (7.4), there was a stark contrast in the secondary structures adopted at low pH (4.0) that could be defined based on the detergent class. Intriguingly, in non-ionic detergent micelles, like DDM [Figure 3B] and β-OG [Figure 3C], the LASV FD maintained a random coil conformation regardless of pH. When we utilized zwitterionic detergent micelles, such as DPC [Figure 3D] and LDAO [Figure 3E], the FD began to adopt a helical conformation with a helical content of 26% and 24%, respectively, at pH 4.0. In LMPG [Figure 3F], an anionic detergent, the LASV FD had the most helical content at a low pH, which was calculated to be 29%. Notably, in neutral bicelles, the LASV FD remained largely random coil regardless of pH and appeared similarly to the non-ionic detergent micelles [Figure 3G]. On the other hand, in acidic bicelles, the LASV FD underwent a conformational change to adopt a helical structure at pH 4.0 with a spectrum similar to DPC and 27% helical content [Figure 3H]. Taken together, it is evident that the LASV FD requires a membrane mimic with charge present to associate with and adopt the helical conformation established to occur at a low pH.

### Membrane mimics with an anionic group provide the strongest resolution of the LASV FD

We next aimed to elucidate the stability and folding of the LASV FD in the most promising membrane mimics, namely LDAO, DPC, LMPG, and acidic bicelles via solution NMR spectroscopy. Additionally, we chose to investigate CHAPS as other zwitterionic detergent micelles appeared to be promising candidates, but we could not effectively examine the secondary structure of the LASV FD by CD spectroscopy. To do this, we quantitated the number of resolved NMR signals, which should match the number of non-proline residues in the protein, and their relative intensities in each spectrum to deduce the optimal detergent for the LASV FD. Since the construct we employed had 36 amino acids, but one proline residue, we would expect that there would be 35 peaks in a ^1^H-^15^N heteronuclear single quantum coherence (HSQC) spectrum. Additionally, some residues, have subtle, but distinct spectral regions in which they will appear, particularly signals from the glycine backbone typically exist at the high edge of the ^15^N dimension, in the 105 – 100 ppm range, whereas serine and threonine residues should have ^15^N chemical shifts of approximately 115 ppm. As such, well-resolved spectra would have approximately 23 peaks in the fingerprint region of the HSQC with ^15^N chemical shifts of 118 – 130 ppm. A ^1^H-^15^N heteronuclear single quantum coherence (HSQC) spectrum revealed dispersed peaks in all five tested membrane mimics, indicative that the LASV FD was well structured [Figure 4]. However, there was sharper spectral resolution of the LASV FD in anionic membrane mimics, namely LMPG and acidic bicelles, over the zwitterionic membrane mimics, chiefly CHAPS, DPC, and LDAO. The DPC spectrum collectively had the poorest resolution out of all the employed detergents with an average peak height and peak volume of 4.43e6 ± 5.05e6 and 8.70e7 ± 1.05e8 ga, accordingly, followed by LDAO (2.24e6 ± 2.96e6 and 4.70e8 ± 5.28e8 ga), and then CHAPS (4.42e6 ± 7.87e6 and 5.53e8 ± 8.71e8 ga) where smaller values are indicative of poorer peak sharpness [Table 1]. Likewise, the average linewidth in both the ^1^H and ^15^N dimensions were smallest in DPC at 44.7 ± 35.5 and 56.5 ± 64.6 Hz, respectively, then LDAO (29.2 ± 38.6 and 97.2 ± 73.2 Hz), and CHAPS (82.2 ± 68.0 and 62.1 ± 111 Hz), suggestive of duller peaks. In contrast, LMPG and acidic bicelles had an average peak height of 2.34e7 ± 2.50e7 and 6.52e6 ± 7.47e6 ga with an average peak volume of 3.63e9 ± 3.58e9 and 1.05e9 ± 1.29e9 ga, correspondingly, suggestive of sharper peaks in LMPG than acidic bicelles. The average linewidth in the ^1^H and ^15^N dimensions were relatively similar in LMPG and acidic bicelles at 113 ± 83.7 and 105 ± 82.5 Hz, accordingly, in the ^1^H dimension and 114 ± 80.9 and 94.0 ± 76.4 Hz for the ^15^N dimension. Taken together, these values culminate to the highest signal-to-noise ratio in LMPG (211 ± 226), followed by acidic bicelles (108 ± 124), LDAO (109 ± 81.0), CHAPS (89.4 ± 74.6), and finally, DPC (46.6 ± 47.3). Nonetheless, despite having the highest signal-to-noise ratio, the LMPG spectrum had only 30 of the 35 expected peaks, including six of the seven expected peaks for serine and threonine residues and five out of five glycine residues. Acidic bicelles, which had a similar resolution to LMPG when the errors of each NMR parameter were considered, had more than the expected peaks, including all of the serine, threonine, and glycine residues. As such, acidic bicelles are the optimal membrane mimic to be employed for further solution NMR structural studies to capture the essence of the LASV FD.

**Table 1.**
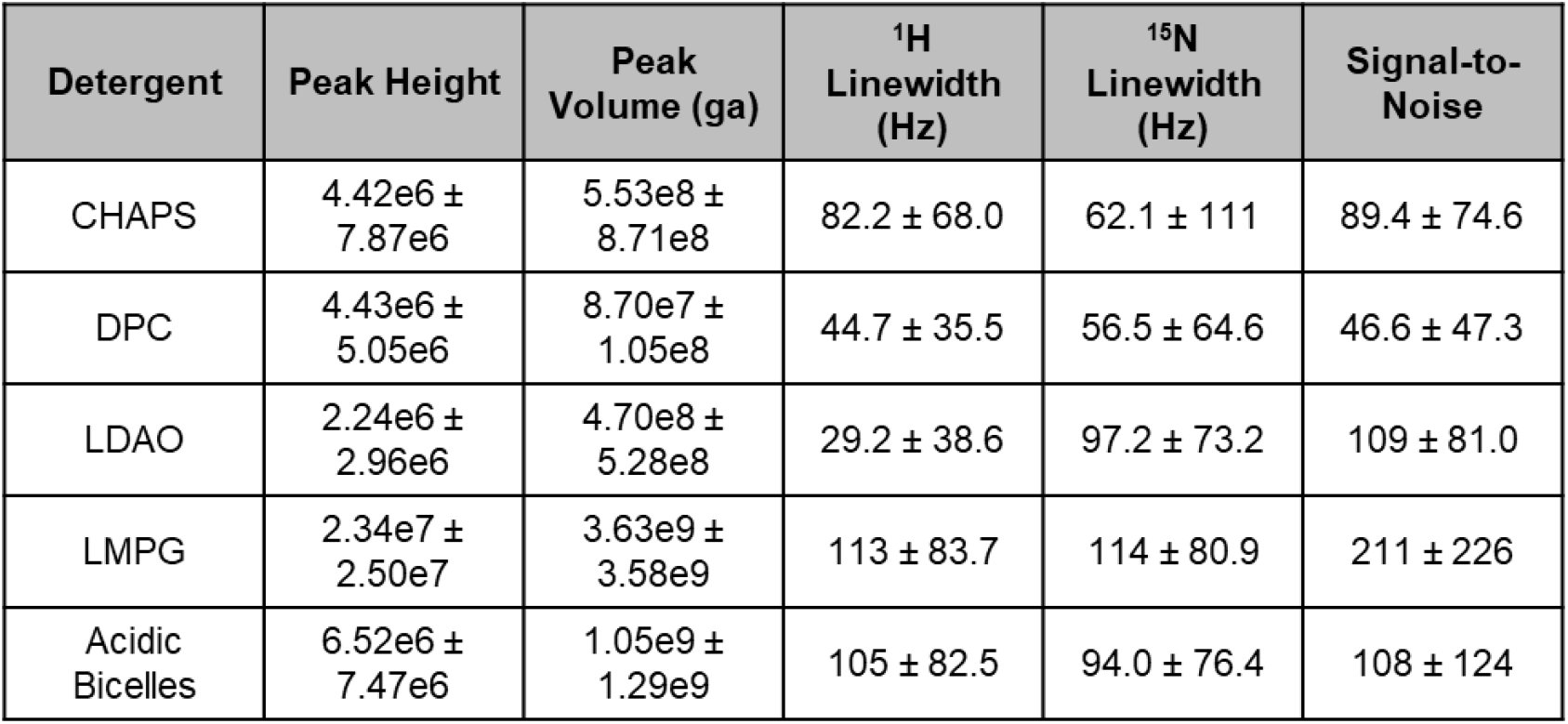
Comparison of the average values for different parameters indicated that the LASV FD had the sharpest resolution in an anionic membrane mimic over zwitterionic membrane mimics. Values calculated using NMRFAM-Sparky.

| Detergent | Peak Height | Peak Volume (ga) | $^1\text{H}$ Linewidth (Hz) | $^{15}\text{N}$ Linewidth (Hz) | Signal-to-Noise |
| --- | --- | --- | --- | --- | --- |
| CHAPS | $4.42\text{e}6 \pm 7.87\text{e}6$ | $5.53\text{e}8 \pm 8.71\text{e}8$ | $82.2 \pm 68.0$ | $62.1 \pm 111$ | $89.4 \pm 74.6$ |
| DPC | $4.43\text{e}6 \pm 5.05\text{e}6$ | $8.70\text{e}7 \pm 1.05\text{e}8$ | $44.7 \pm 35.5$ | $56.5 \pm 64.6$ | $46.6 \pm 47.3$ |
| LDAO | $2.24\text{e}6 \pm 2.96\text{e}6$ | $4.70\text{e}8 \pm 5.28\text{e}8$ | $29.2 \pm 38.6$ | $97.2 \pm 73.2$ | $109 \pm 81.0$ |
| LMPG | $2.34\text{e}7 \pm 2.50\text{e}7$ | $3.63\text{e}9 \pm 3.58\text{e}9$ | $113 \pm 83.7$ | $114 \pm 80.9$ | $211 \pm 226$ |
| Acidic Bicelles | $6.52\text{e}6 \pm 7.47\text{e}6$ | $1.05\text{e}9 \pm 1.29\text{e}9$ | $105 \pm 82.5$ | $94.0 \pm 76.4$ | $108 \pm 124$ |

**Figure 4.**
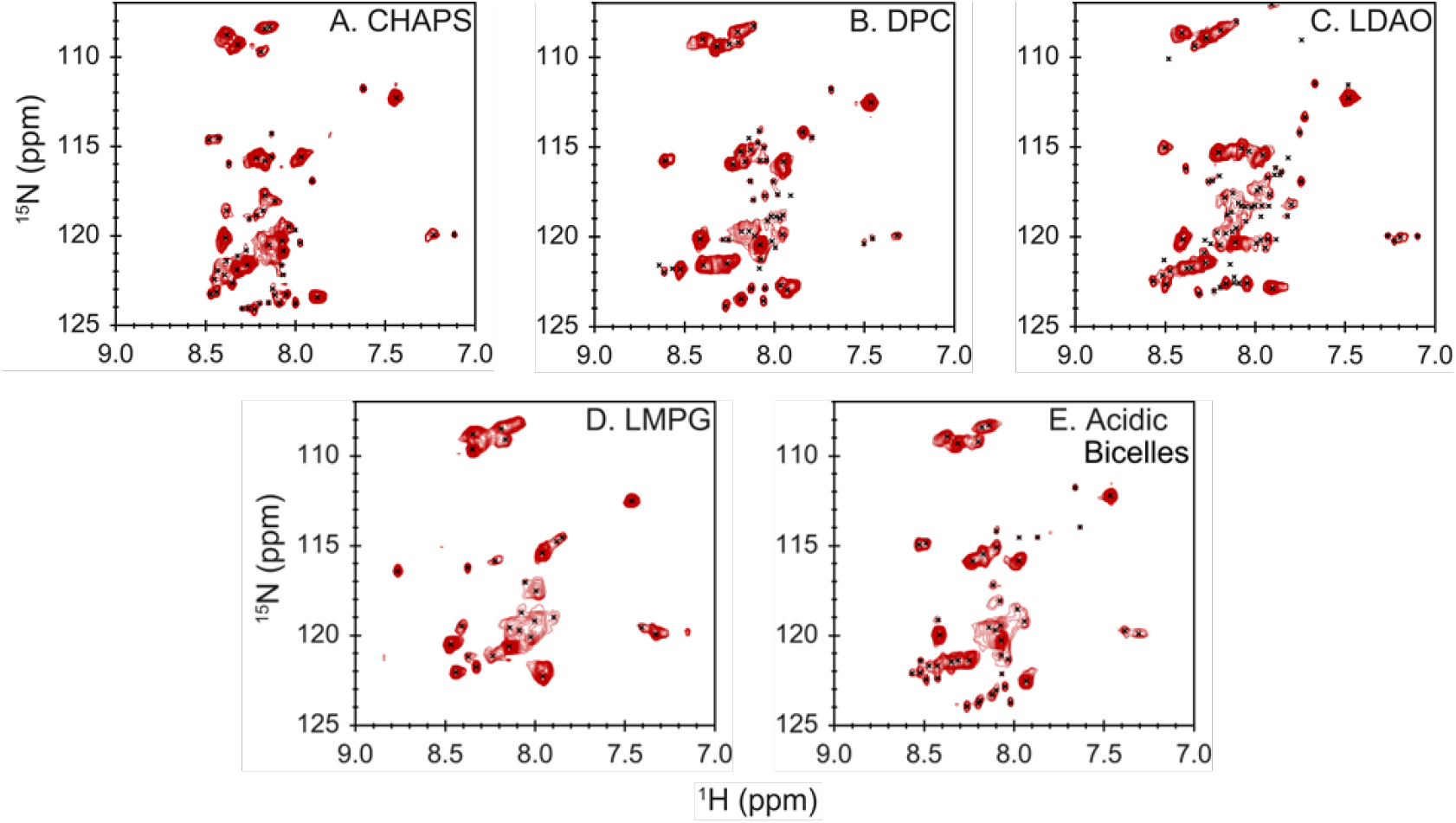
Anionic bicelles provide the best spectral resolution of the LASV FD. In zwitterionic detergent micelles, such as [A] CHAPS, [B] DPC, and [C] LDAO, the signal-to-noise ratio was low, which led to poorly resolved spectra. The LASV FD was well resolved in [D] LMPG and [E] acidic bicelles, which both carry anionic charges, but more well-defined peaks were observed in acidic bicelles.

## Discussion

LASV is a lethal, emerging arenavirus with pandemic potential that has no FDA-approved therapeutic options. The LASV-mediated membrane fusion is initiated by the FD, which makes it an intriguing target for neutralization. Multiple fusion inhibitors have been described for other class one fusion proteins that target their respective FD, such as HIV, influenza, SARS-CoV-2, and EBOV. However, the lack of structural information regarding the LASV FD, particularly after it initiates fusion, has been a limiting factor in designing FD-targeting antibodies. Biophysical characterization of membrane proteins requires elucidation of the most suitable membrane mimic to keep the protein properly folded and in an active state. While the LASV FD is not a traditional membrane protein, it does insert itself into, and thus perturb, the host cell membrane at a low pH [Figure 1], which has made it difficult to elucidate its post-fusion structure. Here, we have demonstrated that the LASV FD has the highest helical content in the presence of anionic membrane mimics, chiefly the negatively charged detergent micelle LMPG and acidic bicelles. We reveal that acidic bicelles are the optimal membrane mimic to move forward with structural studies.

In the literature, the random coil conformation of a class one fusion protein is often regarded as the non-fusogenic form, whereas the helical conformation is the fusogenic form^18, 21, 26, 52-55^. We previously demonstrated that the LASV FD adopted a fusogenic, helical conformation at a pH akin to the lysosomal compartment (pH 4.0)^22^. However, this was accomplished in POPC:POPG SUVs, which are unfavorable for structural studies, particularly NMR spectroscopy, due to their slow molecular tumbling that leads to broad and unresolved spectra^56-58^. Thus, it was necessary to deduce the most optimal membrane mimic for structural studies of the LASV FD. Our CD spectroscopy experiments showed that there is a clear linkage between the class of membrane mimic employed and the ability of the LASV FD to adopt its fusogenic, helical conformation. Intriguingly, the LASV FD had the highest helical content in acidic bicelles, and LMPG, where both have anionic properties [Figure 3]. Notably, the lysosomal compartment has been shown to have a high concentration of anionic lipids, particularly bis(monoacylglycerol)phosphate (BMP)^59-63^. BMP, as well as other anionic lipids, are shown to influence a variety of fusion processes, often through a charged-charged interaction^64-68^. For example, the SARS-CoV-2 FD, which enters the host cell via the late endosomal compartment, has a preference to initiate fusion in the presence of BMP over other anionic lipids^69^. It has also been suggested that BMP serves as an important cofactor to selectively promote LASV-mediated membrane fusion with late endosomes/lysosomes^70^. Therefore, we postulate that the LASV FD prefers to adopt its fusogenic, helical conformation in anionic membrane mimics due to its ability to form similar, favorable interactions that would otherwise occur in the negatively charged lysosomal membrane. This notion can be further supported by our observation that in mild, non-ionic detergent micelles, such as DDM and β-OG, the LASV FD remained in a random coil conformation, regardless of pH, whereas an intermediate helical conformation was adopted in zwitterionic detergent micelles, like LDAO and DPC.

For our NMR studies, the LASV FD had comparable average signal-to-noise ratios, linewidths in the ^1^H and ^15^N dimensions, peak heights, and peak volumes in acidic bicelles and LMPG, which were stronger than CHAPS, DPC, and LDAO, indicative of sharper peaks and better resolution [Figure 4]. Evidently, a negatively charged membrane mimic alone was not enough to stabilize the LASV FD as the number of peaks was greater in acidic bicelles over LMPG at approximately 50 and 30 peaks, correspondingly, where a total of 35 peaks were expected. In the literature, numerous proteins have been shown to have different properties in detergent micelles and bicelles^71-75^. It has been suggested that the inherent ability of detergent micelles to form a spherical, monolayer suppresses the flexibility of membrane proteins, leading to altered properties. On the other hand, bicelles form a non-continuous bilayer that more closely resembles the lipid bilayer and allows a membrane protein to maintain its native conformation. This difference could explain why we observed an excess amount of peaks for the LASV FD in acidic bicelles. In particular, the LASV FD may have high flexibility with several residues adopting multiple conformations on a timescale that can be captured by the NMR, as has been demonstrated with the influenza FD^19, 75-77^. In LMPG, however, the NMR spectra of the LASV FD appeared to reflect the average signal of a conformation, rather than capturing multiple conformational states, likely due to a suppression of dynamics correlated with detergent micelles. Moreover, while it has previously been suggested that acidic bicelles have issues with stability at a pH below 5.5^78^, we present evidence that they are stable for at least a week at pH 4.0, which would provide adequate time to perform the 3D experiments necessary via solution NMR spectroscopy to elucidate the post-fusion structure [Figure 2]. Therefore, not only does the LASV FD require the presence of a negatively charged membrane mimic, but we also provide evidence that a lipid bilayer was required for suitable spectral resolution for structural studies, such as acidic bicelles.

In summary, our data indicates that acidic bicelles are the optimal membrane mimic for structural studies of the LASV FD. We have illustrated that an anionic charge alone is insufficient to stabilize the LASV FD and an acidic, bilayer-like system must be employed to capture the native structure. Overall, these findings will allow for structural elucidation of the LASV FD to understand how it perturbs the host cell membrane. Revealing the structure of the LASV FD has the potential to provide critical information for designing therapeutic agents to target the FD and prevent LASV infection.

## Author Contributions

H.N.P and J.L. designed the experiments. H.N.P performed the experiments and analyzed the data. H.N.P and J.L prepared the manuscript.

## Acknowledgements

We would like to thank all lab members for their assistance in editing this manuscript.

